# Temperature predicts the maximum tree-species richness and water and frost shape the residual variation

**DOI:** 10.1101/836338

**Authors:** Ricardo Segovia

## Abstract

The kinetic hypothesis of biodiversity proposes that temperature is the main driver of variation in species richness, given its exponential effect on biological activity and, potentially, on rates of diversification. However, limited support for this hypothesis has been found to date. I tested the fit of this model on the variation of tree-species richness along a continuous latitudinal gradient in the Americas. I found that the kinetic hypothesis accurately predicts the upper bound of the relationship between the inverse of mean annual temperature (1/kK) and the natural logarithm of species richness, at a broad scale. In addition, I found that water availability and the number of days with freezing temperatures organize a part of the residual variation of the upper bound model. The finding of the model fitting on the upper bound rather than on the mean values suggest that the kinetic hypothesis is modeling the variation of the potential maximum species richness per unit of temperature. Likewise, the distribution of the residuals of the upper bound model in function of the number of days with freezing temperatures suggest the importance of environmental thresholds rather than gradual variation driving the observable variation in species richness.

## Introduction

The geographic variation in species richness over broad spatial scales is probably the most intriguing pattern of biodiversity. While correlations between species richness and environmental variables have been widely documented (Adams and Woodward, 1989; Currie, Mittelbach, et al., 2004; Currie and Paquin, 1987; Field et al., 2005; Francis and Currie, 2003; Hawkins, Porter, et al., 2003; Jetz and Rahbek, 2002; Kreft and Jetz, 2007; Mutke and Barthlott, 2005; Mutke, Kier, et al., 2001; O’Brien, 1993, 1998; Stephenson, 1990; Wright, 1983), a general law or even a single, coherent explanation for these correlations is still elusive (Brown, 2014; Gaston, 2000; Rohde, 1992). The most formal and testable hypothesis to mechanistically relate species richness to environmental parameters is the kinetic model, which forms part of the framework of the “Metabolic Theory of Ecology” (Brown, 2014). Under this hypothesis, temperature would be the main driver of variation in species richness, given its effect on the activation of enzymatic reactions, the rates of energy flux and thus, potentially, on rates of ecological interaction and diversification (Allen, Brown, et al., 2002; Brown, 2014; Brown et al., 2004; Rohde, 1992). Therefore, a predictive model derived from the exponential Boltzmann temperature relationship has been proposed and tested across different taxa (Allen, Brown, et al., 2002; Brown et al., 2004). However, subsequent studies at global (Kreft and Jetz, 2007) and regional (Algar et al., 2007; Hawkins, Albuquerque, et al., 2007) scales show no support for two of the main predictions of the model: 1) a linear relationship between species richness and temperature (Allen, Brown, et al., 2002; Allen, Gillooly, et al., 2007; Brown et al., 2004), and 2) a negative slope of ~ 0.65 between the inverse of temperature (1/*k*K) and the natural logarithm of species richness (Brown et al., 2004).

To predict species richness gradients, the kinetic model connects two previously recognized relationships: the temperature dependence of the metabolic rate (Gillooly et al., 2001) and the energetic-equivalence rule of population energy use (Enquist, Brown, et al., 1998). Temperature controls metabolism through its effects on rates of biochemical reactions. Reaction kinetics vary exponentially with temperature according to the Boltzmann’s factor *e^−Ei/kT^*, where T is the absolute temperature (in degrees K), *E* is the activation energy, and *k* is Boltzmann’s constant (Gillooly et al., 2001). The increase in metabolic rate results in turn in higher energy use per individual, but the population energy use should remain approximately constant (Enquist, Brown, et al., 1998). Thus, fewer individuals per population are expected towards warmer environments (Brown et al., 2004). Considering that the total number of individuals in a community is largely independent of latitudinal gradients in temperature when no other environmental constraints are influencing it (i.e., water deficit) (Currie, Mittelbach, et al., 2004; Enquist and Niklas, 2001), the result is higher species richness in communities from warmer environments (Allen, Brown, et al., 2002). Moreover, the kinetic model suggests that in addition of the direct effect of energy use increasing, coevolutionary interactions associated with species coexistence generate and maintain even higher biodiversity towards warmer environments (Brown, 2014).

In addition to temperature, the variation in species richness has also been related to water availability, often measured as mean annual precipitation (Currie, 1991; Wright, 1983). Therefore, the effect of water availability could explain the failure of previous studies to support the temperature dependence of species richness across different environmental arrangements (Currie, 2007). Currie (2007) hypothesized that the model proposed by Allen *et al.* (2002) (Allen, Brown, et al., 2002) only would fit on the upper bound of the relationship between species richness and mean annual temperature (MAT) in plants, but cannot explain the species richness variation on its mean values. Allen *et al.* (2007) agree with this, and suggest that the residual variation below the proposed upper bound fit would be associated with variation in the total number of individuals in a community due, mainly, to water deficit. Indeed, statistical models show that forests can reach their maximum species richness when high annual energy input is combined with high water availability throughout the year (Kreft and Jetz, 2007). However, limits rather than mean temperature also could explain part of the residual variation of the kinetic model. The metabolism-temperature dependence is valid within the limited range of “biologically relevant” temperatures between approximately 0° and 40°C. Near 0°C, metabolic reactions cease due to the phase transition associated with freezing water (Brown et al., 2004; Gillooly et al., 2001). Thus, events of freezing temperatures can interrupt the general effect of environmental temperature on biochemical reactions and energy use, modifying the expected species richness beyond the effect of mean values (i.e., MAT).

The original proposal of the kinetic model and the posterior empirical approaches have only tested the central tendency of the variation in species richness (Algar et al., 2007; Allen, Brown, et al., 2002; Brown et al., 2004; Hawkins, Diniz-Filho, et al., 2007; Hawkins, Albuquerque, et al., 2007; Kreft and Jetz, 2007), rather than the upper bound of the relationship. In order to evaluate the predictions of the kinetic model around the upper bound of the temperature dependent variation in species richness, I analyzed 10,721 inventories of tree species across a broad, continuous latitudinal gradient in the Americas (between 49°N and 55°S in North, Central and South America). I tested the predictions of linearity and a negative slope (~ 0.65) using regression models throughout the upper quantiles of species richness variation. To test the linearity of the relationship between the inverse of temperature (1/*k*K) and the upper quantiles of the natural logarithm of species richness, I assessed if segmented models, that allow for breakpoints in the slope at some point over the range of the explanatory variable, provided better statistical fits than linear models. Thus, I compared the slopes of linear versus segmented models above the mean of the relationship (55%, 65%, 75%, 85%, 95% quantiles). In addition, I explored the influence of the previously hypothesized environmental drivers (i.e., climatic water deficit and freezing temperatures) on the variation of the residuals of quantile regressions.

## Results

As increasingly higher quantiles are examined (i.e. moving from 55% quantiles to 95% quantiles), the statistical relationship between temperature (1/*k*K) and the natural logarithm of species richness becomes stronger (Table 1). In fact, the values for both linear and segmented models reach over R1 = 0.7 at 95% quantile (see R1 statistic, Table 1). Although segmented models are better fitted in all of the quantiles analyzed (see SIC, Table 1), both of the slopes in the segmented models towards the upper bound are not significantly different from each other (Table 1, p-values greater to 0.05 to the test of the null hypothesis, slope 1 = slope 2). Furthermore, towards the very upper bound (quantile 85% and 95%) both slopes are negatives, while in 55%, 65% and 75%, the first slope is positive. Therefore, a linear negative relationship towards the upper bound, where both models fit the better, cannot be rejected (Table 1, Fig. 1).

**Table 1.**
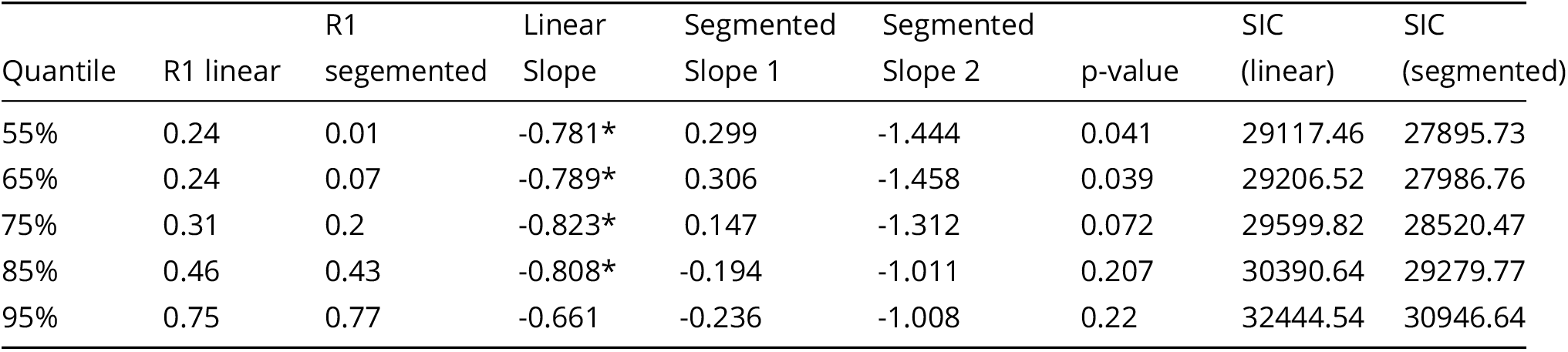
Quantile models across the upper bound of the relationship between the inverse of temperature (1/*k*K) and the natural logarithm of species richness. R1 statistic is provided as a measure of the goodness of fit of the different models and the Schwartz’s Information Criteria (SIC) as a relative quality measurement for linear and segmented models (for details, see Material and Methods). Slope values for linear and segmented quantile regressions (slope 1 and slope 2) are also provided. Asterisks in values for linear slopes show those slopes that statistically differ from the theoretical values of −0.65. Finally, the p-value column means the results of the *t* tests comparing if both slopes in the segmented models are significantly different (p-values < 0.05).

**Fig. 1.**
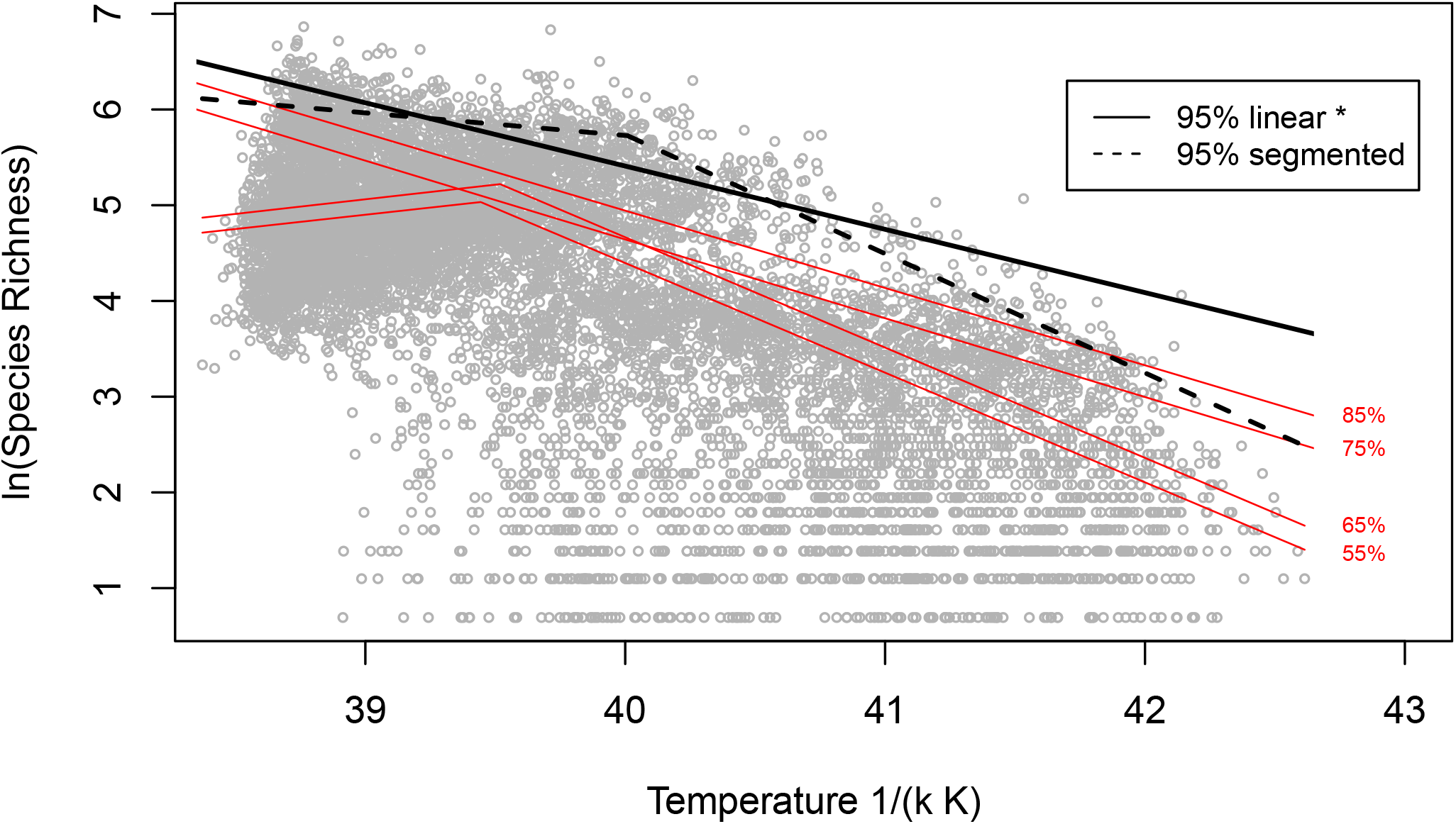
Effects of the inverse of the Mean Annual Temperature (MAT) on log-transformed species richness of trees in the Americas. Black lines represent the model fit at 95% quantile for a linear regression (continuous) and for a one-breakpoint segmented regression (dashed). The red lines represents the best supported model (linear or segmented) in 55%, 65%, 75% and 85% quantiles. Breakpoint models were considered “best supported” only if the slopes of both of the segments statistically differ from each other.

I found that the negative linear slope of ~ 0.65, proposed by Brown *et al.* (2004) (Brown et al., 2004) is reached around the 95% quantile (Table 1, Fig. 1, p-value=0.42 to test the hypothesis, slope in 95% quantile = −0.65). Instead, slopes values for those linear model in lower quantiles statistically differ from the theoretical values expected by the kinetic model (Table 1, Fig. 1, p-value < 0.005 to test the hypothesis, slopes = −0.65, for 55%, 65%, 75% and 85% quantiles). On the other hand, segmented models are far from the predicted range of slopes put forth by Brown *et al.* (2004) (Table 1).

The temperature dependence of species richness at the upper bound analyzed across biomes shows that the linear slopes are mostly different than the expected by the model (Fig. 2). Slopes for the 95% quantile model from tropical biomes are farther away of the expected values than those from temperate biomes (Fig. 2). Likewise, dry biomes present the most different slopes according to the expected under the kinetic model. Slopes in dry biomes tend to be positive rather than negative, i.e., they show a greater potential species richness towards relatively colder areas within “Tropical & Subtropical Grassland, Savannas Shrublands”, “Tropical & Subtropical Dry Boroadleaf Forests” and “Mediterranean Forests, Woodlands and Scrub” (Fig. 2). The closest slopes to the expected by the kinetic model are found in “Temperate Grasslands, Savannas & Shrublands” and “Temperate Conifer Forests” (Fig. 2).

**Fig.2.**
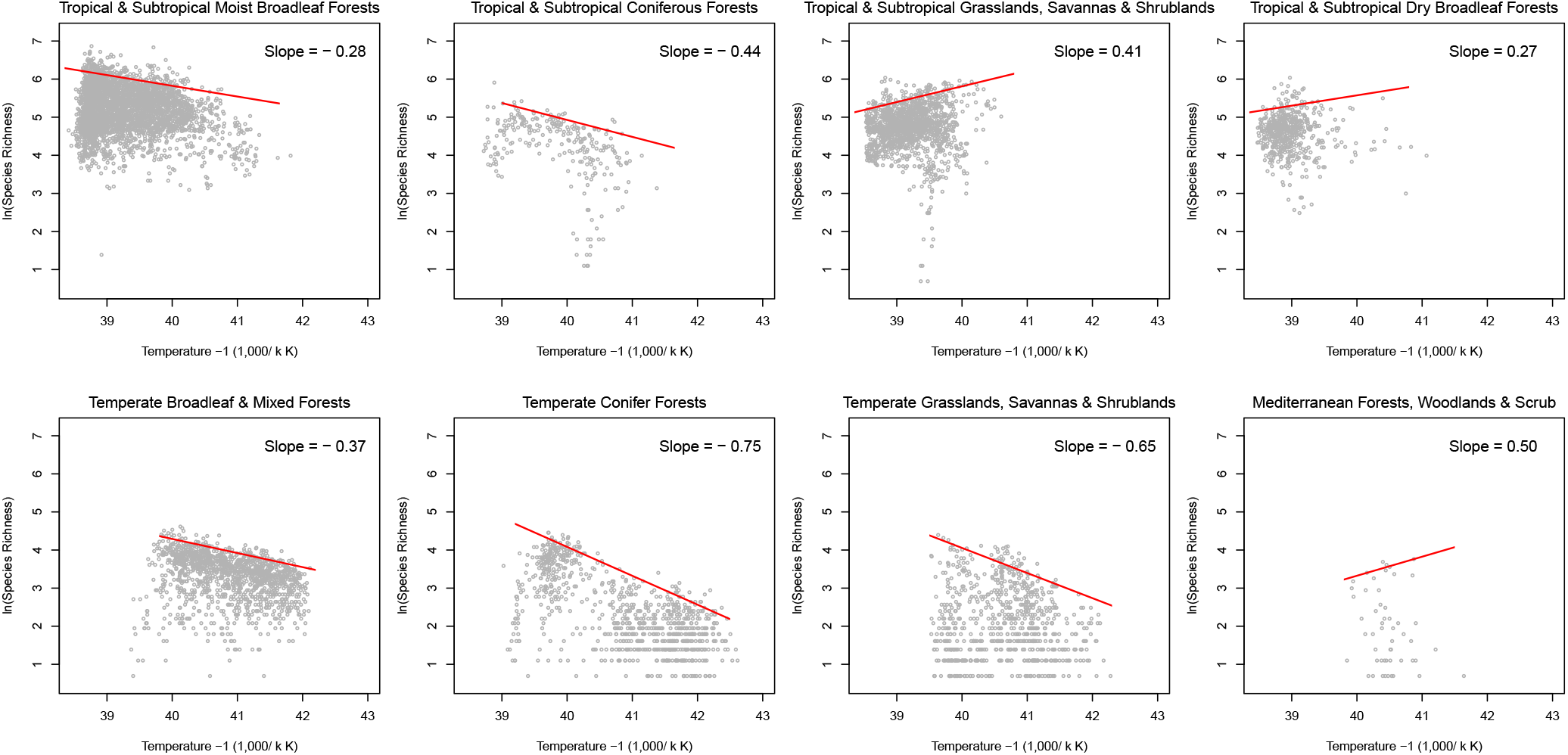
Quantile 95% model across the main biomes in the Americas.

In a general view, the spatial distribution of the residual variation in the upper bound model shows that sites with low species richness are farther away from the theoretical predictions than species-rich sites (Fig. 3). Sites with residuals lower than −1.5 tend to be concentrated preferentially in the northern and southern extratropics, subtropics and dry tropics (Fig. 3). In a biome-organized perspective, the residual variation in the 95% quantile model is more negative on average in temperate than in tropical biomes (Fig. 4). Likewise, sites from tropical wet biomes tend to be closer to the upper bound model than sites from tropical dry, subtropical and extratropical environments (Fig. 4).

**Fig.3.**
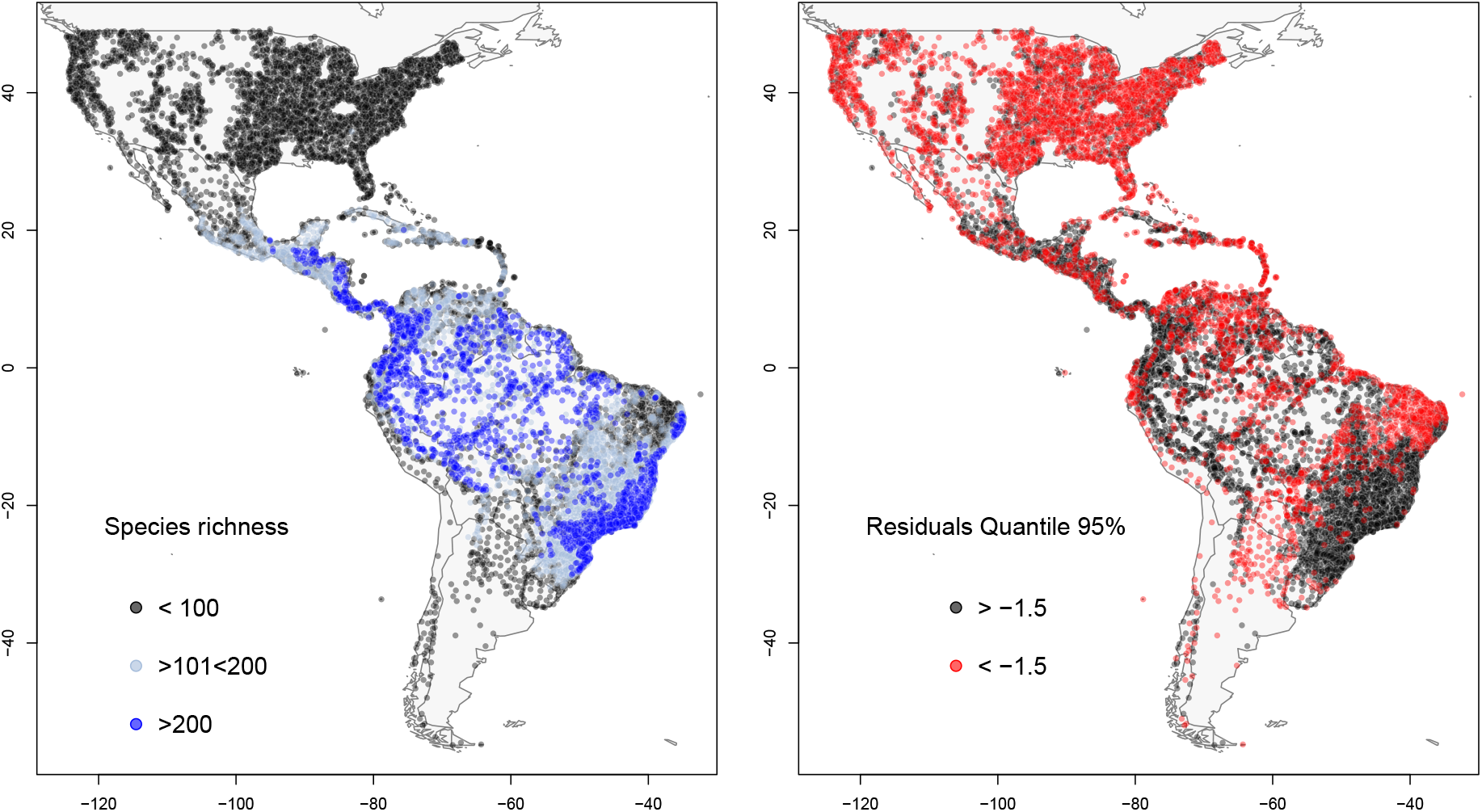
Geographic distribution of the species richness variation and the residual variation of the quantile 95% model. A) Species richness across the 10,721 tree assemblages. Black dots show those assemblages with less than 100 species. Light-blue dots show those assemblages with a number of species between 101 and 200. And, blue dots show assemblages with more than 201 species. B) Residual variation from the upper bound model for the kinetic hypothesis. Black dots show those assemblages with more negative residual variation, and red dots those assemblages with a less negative residual variation.

**Fig.4.**
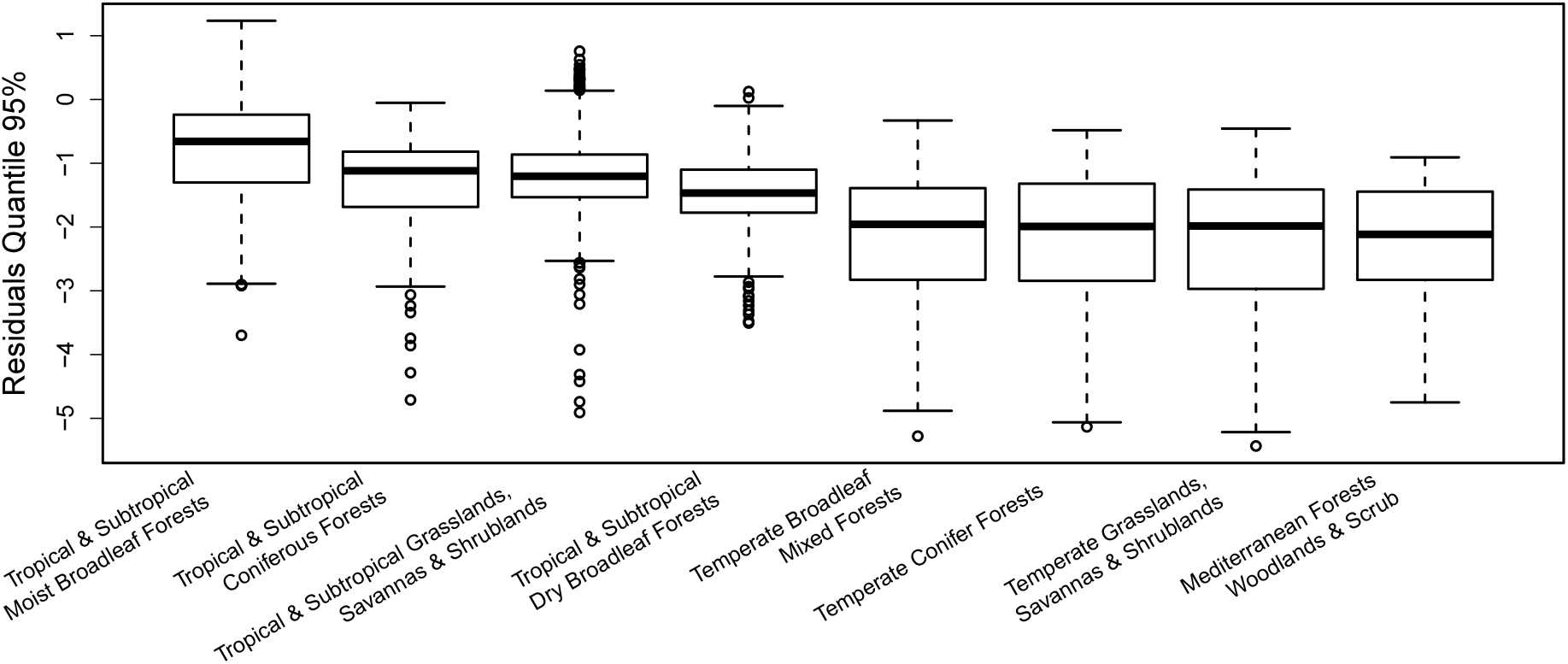
Residual variation of the quantile 95% model across the main biomes in the Americas. Biome partition of the dataset is based on the ecorregions and biomes described by Olson *et al.* (2001) (Olson et al., 2001).

Water availability and presence of freezing temperatures affect the residual variation around the upper bound of the kinetic model for the temperature dependence of species richness (Fig. 5). Residual variation tends to increase (i.e. more negative residuals) towards environments with higher Climatic Water Deficit, indicated by more negative values (Fig. 5a), and with greater exposure to freezing temperatures (Fig. 5b). While the effect of water deficit shows a relatively gradual change, the effect of freezing temperatures suggests a threshold effect (Fig. 5). Those sites with more than ~ 1,000 days with freezing temperatures during the last 117 years (i.e., greater than ~ 10 days on average per year) tend to be farther away of the predicted model for the 95% quantile than those sites with less than 10 days per year subject to freezing temperatures (Fig. 5b).

**Fig.5.**
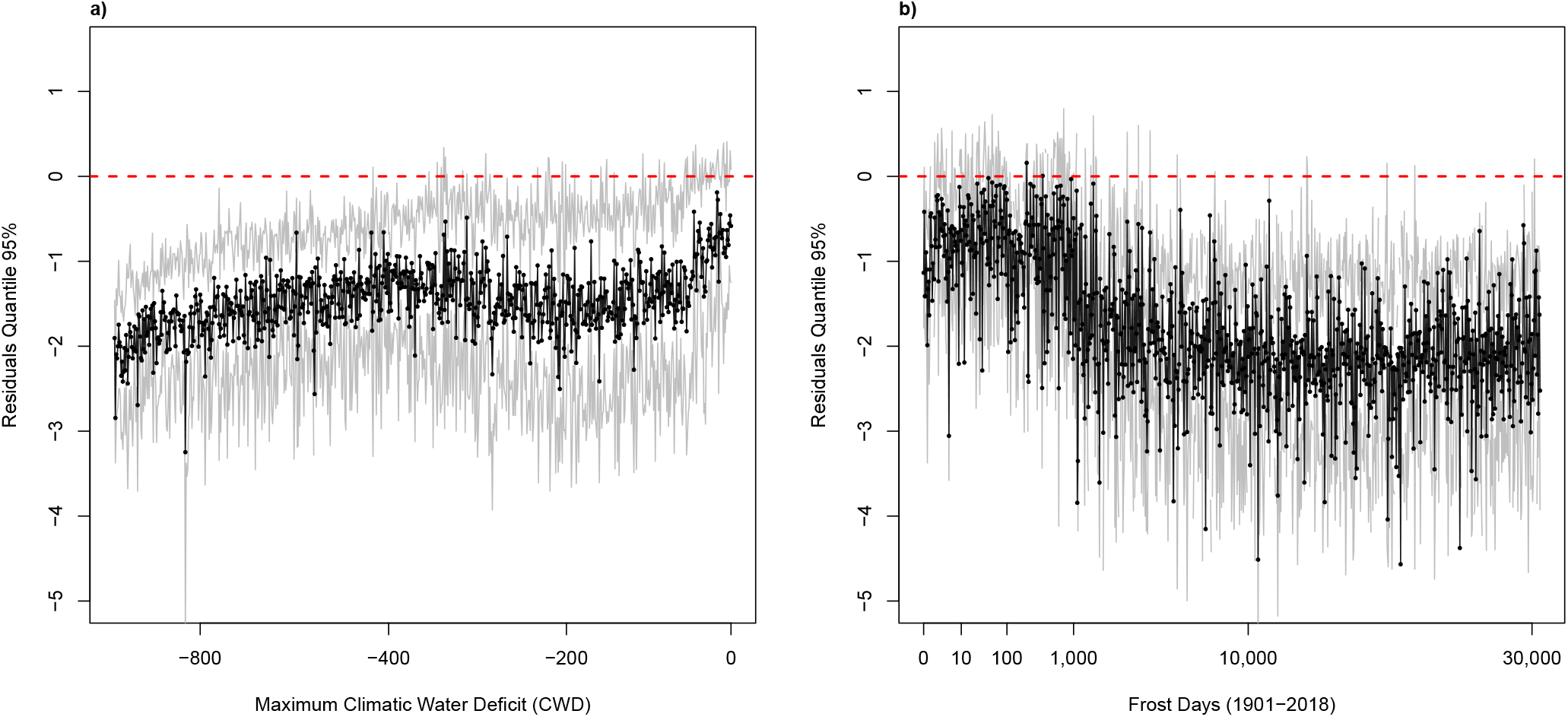
Distribution of the residuals according to the 95% quantile fit along two limiting species richness environmental variables: A) the Maximum Climatological Water Deficit, and B) the Number of Frost Days between 1901 and 2018. Black lines connect the points showing the mean values for each ordered category and gray lines connect the points for standard deviation.

## Discussion

These results show that the kinetic model that mechanistically relate the variation in species richness with environmental temperature apply to the upper bound of the relationship in a broad scale. Both of the central predictions of the model (i.e., a slope a) linear of b) ~ −0.65) are found at the 95% quantile of the regression (Table 1, Fig. 1). Two environmental drivers impose constraints to the assemblages in reaching the potential maximum species richness. The greater the water deficit and the number of days with freezing temperatures in a year, the bigger the residual variation of the upper bound model (Fig. 5).

The kinetic hypothesis (Allen, Brown, et al., 2002; Brown et al., 2004) models a limit for the maximum species richness per unit of temperature rather than variation in the mean values. Although later testing of the kinetic hypothesis focused on the temperature-dependent variation in species richness across different environmental arrangements, the original proposal specified that the model works only if other environmental variables (e.g., water availability) are accounted for (Allen, Brown, et al., 2002; Allen, Gillooly, et al., 2007). In concordance with the original proposal, the results presented here suggest that under the kinetic model, temperature determines an upper bound of variation in species richness (Fig. 1), and that other variables may prevent local assemblages to reach this potential diversity (Fig. 5). In other words, temperature would determine an environmental carrying capacities rather than to drive the observable variation in species richness, and the kinetic hypothesis is actually modeling the variation of such limits. Thus, under a bounded framework for diversity, the environmental temperature affecting productivity or energy availability should be interpreted as an ecological constraint placing limits to the potential maximum species richness (see (Brodie, 2019; Cornell, 2013)).

Considering temperature as an ecological driver of variation in the environmental carrying capacity should have implications on evolutionary biology and other fields of ecology. For example, some broadly used assumptions to estimate diversification rates should be reconsidered. Usually, phylogenetic studies use the ratio between clade diversity and clade age to estimate diversification rates and do not consider environmental constraints to diversification (e.g., (Magallon and Sanderson, 2001; Nee, 2001; Phillimore et al., 2006)). However, if species richness is environmentally constrained, this ratio will be misleading since diversity is no longer dependent only on time and diversification rate (Cornell, 2013). In other examples, community assemblage under a bounded or an unbounded framework should have different rules. If diversity is bounded by ecological constraints, or if ecological constraints are at least slowing diversification, then the asymptote of the diversification rate should be correlated with a decrease in niche space (Cornell, 2013). Thus, a theoretical trajectory to the niche space filling associated with an increasing in both taxonomic and traits diversity should be possible to hypothesize.

The upper bound fit for the kinetic model works at global (i.e., a complete latitudinal gradient in the Americas) but not at the level of biomes. As the results presented here show, a biome-partitioned analysis of the slopes at the upper bound do not fit with the theoretical predictions as a generality (Fig. 2). Therefore, the expectations of a kinetic model refocused in the upper bound should be tested considering all regions/biomes under the same model; or, in regional scales, under gradients of temperature with other variables being controlled, as the original proposal suggests (Allen, Brown, et al., 2002). Biomes are evolutionary modules environmentally constrained, with functional identities (Mucina, 2019). Thus, different environmental arrangements are expected to drive the observable variation in diversity within biomes. Instead, the potential maximum species richness expected per temperature is a general expectation and should be tested in broader scales to avoid the regional effect of evolutionary associations between a particular biota and the particular environmental arrangement constraining the distribution of its diversity.

The slopes of temperate biomes fit better than tropical ones to the predictions of the kinetic model for the upper bound (Fig.2). Indeed, the variation of species richness at a global scale is not stationary (Hawkins, Porter, et al., 2003), and temperature is a better predictor towards high latitudes (i.e., in temperate biomes) (Currie, Mittelbach, et al., 2004; Field et al., 2005; Hawkins, Porter, et al., 2003; Kreft and Jetz, 2007). Instead, the species richness variation at low latitudes (i.e., in tropical biomes) would be best predicted by water-availability variables (e.g., actual evapotranspiration and annual rainfall) (Currie, Mittelbach, et al., 2004; Field et al., 2005; Hawkins, Porter, et al., 2003; Kreft and Jetz, 2007), or biotic factors (MJ Donoghue, 2008; Ricklefs, 2004; JJ Wiens, 2011). Under a refocused kinetic model as a predictor of the maximum species richness, those biotic and abiotic factors driving the observable species richness at low latitudes should be interpreted as constraints for the assemblages to reach their potential diversity. A non-stationary pattern of environmental drivers should also be expected to drive the observable variation in low latitudes. For example, different variables should explain the species richness variation along humidity or elevational gradients. In temperate biomes, temperature would describe better the observable variation in tree-species richness (Currie, Mittelbach, et al., 2004; Field et al., 2005; Hawkins, Porter, et al., 2003; Kreft and Jetz, 2007) because the axis of variation in precipitation is shorter than in tropical environments. For example, trees, as a form of life, cannot inhabit dry and cold environments.

Although the slopes of the upper bound kinetic model fit better in temperate biomes (Fig. 2), the distribution of the residuals shows that they are farther away of the model than tropical (wet and dry) ones (Figs. 3, 4). Therefore, the species richness in temperate biomes follow better the theoretical slope predicted according the variation in environmental temperature, but they are homogeneously constrained by other factors. In other words, a climatic feature (beyond of the relative low environmental temperature) of temperate biomes is preventing the assemblages in reaching the maximum potential species richness. The presence of freezing temperatures in a normal year is a feature of temperate biomes and its effect on the variation of species richness has been largely recognized, even by Alexander Von Humboldt (Hawkins, 2001). As the results presented here show, the distribution of the residual variation driven by the number of days with freezing temperatures shows a pattern like a threshold rather than a gradient (Fig. 5b). While assemblages subject to less than ~ 10 days per year with freezing temperatures are close to the model, those assemblages beyond that threshold are homogeneously farther away from the upper bound model (Fig. 5b). The effect of freezing temperature on biodiversity has been detected also in an evolutionary scale, since the best environmental predictor for the evolutionary turnover of tree lineages between tropical and extratropical biotas is the presence or absence of freezing temperatures in a normal year (Segovia et al., 2020). Although a putative mechanism for freezing temperatures preventing the assemblages to reach the potential species richness expected by environmental temperature is the cessation of metabolic reactions due to the phase transition associated with freezing water (Brown et al., 2004; Gillooly et al., 2001), its effect on the evolutionary scale may be related with eco-evolutionary processes that prevent the dispersion and diversification into “harsh”, no tropical (i.e., dry and/or freezing) environments (Currie, Mittelbach, et al., 2004; J Wiens and M Donoghue, 2004).

Freezing temperatures and water deficit have been considered among the primary forces shaping plant evolution by acting directly on hydraulic traits (Maherali et al., 2004; Zanne et al., 2014). Likewise, both of these environmental conditions characterize “harsh”, or difficult to colonize, environments, and thus, would explain a decrease in species richness (J Wiens and M Donoghue, 2004). However, their influence on the variation of the observable species richness work in different ways. While freezing temperatures would slow down diversification by ceasing the temperature-dependent metabolic rate, water deficit may constrain the potential species richness by dropping the total number of individuals in a community (Allen, Brown, et al., 2002; Allen, Gillooly, et al., 2007; Enquist and Niklas, 2001; Gaston, 2000). The results presented here show that the slopes in tropical dry biomes are positive rather than negative (Fig. 2). This trend of potential maximum species richness increasing to relatively colder sites within tropical dry environments is expectable because temperature and precipitation may interact in determining the species richness of water-limited biomes. Actually, the increment of water climatic deficit (i.e., the interaction between temperature and precipitation) drives an increase in the residual variation (Fig. 5). Therefore, future works should be careful in separating the variation in species richness associated with changes in the total number of individuals per unit of area from those with no changes on this variable.

Changing the focus from the central tendency towards the limits allows for clarifying that the metabolic effect of environmental temperature works on the maximum species richness and, therefore, to conclude that the kinetic hypothesis described a model of the potential maximum species richness variation or the environmental carrying capacity. Instead, the observed variation of species richness is controlled by secondary environmental arrangements in different ways, suggesting that its effect can be spatial or geographically not stationary. The structure of wet-tropics species-rich versus dry-tropics and extratropics species-poor can be driven by processes such different as changes in abundances or evolutionary constraints. A perspective that considers that temperature determines an upper bound and other variables can influence the observable species richness through different mechanism should allow a deeper understanding of the relationship between environments and biodiversity.

## Methods

### Dataset

The tree assemblages dataset was derived by combining the NeoTropTree (NTT; http://neotroptree.info) database (Oliveira-Filho, 2014) and the Forest Inventory and Analysis (FIA) Program of the U.S. Forest Service (Burrill et al., 2018), accessed on March, 2019 via the BIEN package (Maitner et al., 2018) for the R Statistical Environment (R Core Team, 2018). I excluded from the FIA database any plot that had less than two species. The inventories in the NTT database are defined by a single vegetation type within a circular area with a 5 km radius and contains records of tree and tree-like species, i.e., freestanding plants with stems that can reach over 3m in height (see http://www.neotroptree.info and Silva de Miranda *et al.* (2018) (Silva de Miranda et al., 2018) for details). Each FIA plot samples trees that are > 12.7 cm diameter at breast height (dbh) in four subplots (each being 168.3 m2) that are 36.6 m apart. I aggregated plots from the FIA dataset in a 10 km diameter space, to parallel the spatial structure of the NTT database. This procedure produced a total dataset of 10,721 tree assemblages distributed across major environmental and geographic gradients in the Americas. Taxonomic nomenclature was made consistent by querying species names against The Plant List (http://www.theplantlist.org). Mean annual temperature was extracted from the Worldclim dataset (Hijmans et al., 2005), the number of frost days per year was extracted from the Climatic Research Unit’s (CRU) (Harris et al., 2020), and Maximum Climatological Water Deficit (CWD) from Chave *et al.* 2014 (Chave et al., 2014) for the coordinates assigned to each tree assemblage.

### Statistical Analyses

#### Kinetic hypothesis testing

In order to explore the fit of the relationship between the inverse of temperature (1/*k*K) and the natural logarithm of species richness towards the upper bound, I used Quantile Regressions (Koenker and Bassett Jr, 1978) above the mean linear regression (50%). As quantile regression is insensitive to heteroscedastic errors and dependent variable outliers (Koenker and Bassett Jr, 1978), it is useful to explore the prediction of the kinetic hypothesis above the mean values. Indeed, quantile regressions have been previously described as particularly useful in exploratory and inferential analyses concerning limiting factors in ecology (Cade et al., 1999), which makes it useful to study the relationship of species richness and temperature towards the upper bound. Thus, I tested the two main predictions derived from the kinetic hypothesis: linearity and a negative slope of −0.65.

#### Linearity

I fitted linear Quantile Regression and one-breakpoint segmented Quantile Regression in 55%, 65%, 75%, 85% and 95% quantiles, with “quantreg” (Koenker, 2018) and “segmented” (Muggeo, 2008) packages in R (R Core Team, 2018). I measured the goodnessof-fit (R1) of each quantile models by estimating the ratio between the sum of absolute deviations in the fully parameterized models and the sum of absolute deviations in the null (non-conditional) quantile model, following (Koenker and Machado, 1999). R1 constitutes a local measure of goodness of fit for a particular quantile rather than a global measure of goodness of fit over the entire conditional distribution, like R2 (Koenker and Machado, 1999). This estimation was done using the function *goodfit* from “WRTDStidal” R package.

In order to define if a linear or a segmented fit is better in each quantile, I used two approaches. First, I compared the models by the Schwartz’s Information Criterion (SIC). Second, following Hawkins et al. (2007a) (Hawkins, Albuquerque, et al., 2007), I tested if both slopes in the segmented model are different from each other using a *t* test. If the two slopes were not significantly different (P > 0.05), the relationship between rescaled temperature and ln-transformed richness was classified as being linear on the quantile, whereas data sets with significantly different slopes were classified as being nonlinear.

#### Slope

Brown *et al.* (2004) (Brown et al., 2004) and Allen *et al.* (2007) (Allen, Gillooly, et al., 2007) argued that slopes of richness–temperature regressions should fall near ~ −0.65. This value is different than the proposed by Allen et al. 2002 (Allen, Brown, et al., 2002) (−0.9) due to later changes in the rescaling transformation. While Allen *et al.* (2002) used as transformation 1000/K, where K is kelvins, Brown *et al.* (2004) and Allen *et al.* (2007) used 1/(*k*K), where *k* is the Boltzman’s constant. In order to test this prediction towards the upper bound of the relationship, I simply compared the slopes of all the linear Quantile Regressions, and each segments in segmented Quantile Regressions. For a more specific approach, I tested if slopes in linear models statistically differ from the theoretical values of −0.65 using a *t* test.

Also, I tested the slopes for the predictions through biomes by splitting the dataset according Olson *et al.* (2001) ecorregions and biomes (Olson et al., 2001). I estimated the slopes for the 95% quantile to tests the predictied slope in an upper bound model to the kinetic hypothesis for each of the eight main biomes in the Americas.

#### Residuals Variation

As I found the 95% quantile regression is the best model to fit the kinetic model for species richness, I explored the spatial distribution and the environmental determinants of the residual variation at this quantile. In order to show the geographic distribution of the residual variation and its relationship with the species richnees, I grouped the residuals in higher than −1.5 (i.e. small residual variation) and lower than −1.5 (i.e. big residual variation). Also I grouped the assemblages by species richness: less than 100 species; between 101 and 200 species; and more than 201 species. Likewise, in order to show the variation in the distribution of residuals across biomes, I plotted the residual variation in a boxplot organized by biomes from Olson *et al.* (2001) (Olson et al., 2001).

Finally, I also tested the variation of the residuals with the variation of Maximum Climatological Water Deficit (CWD) from Chave et al. (2014) (Chave et al., 2014) because Allen *et al.* (2007) (Allen, Gillooly, et al., 2007) hypothesized that the residual variation on an upper bound fit should be associated with water availability. Alternatively, I evaluated the effect of cumulative frost days from 1901 until 2018, from the Climatic Research Unit’s (CRU) (Harris et al., 2020) on the residual variation because the frost line has been proposed as an environmental threshold for taxonomic, functional, and evolutionary diversity on trees and, therefore, I hypothesized that it could also affect the relationship between species richness and mean annual temperature. In order to plot the distribution of residuals across the continuous environmental variables, I use the function discretize() in the “arules” package (Hahsler et al., 2019) with the “cluster” method. Thus, I calculated a mean and a standard deviation across the ordered categories in both of the variables.

## Data accessibility

Data table and code (*.Rmd*) needed to re-do the analyses and figures are available online on Zenodo (https://doi.org/10.5281/zenodo.4606783).

## Acknowledgements

Version 4 of this preprint has been peer-reviewed and recommended by Peer Community In Ecology (https://doi.org/10.24072/pci.ecology.100075). I thanks Dr. Kyle Dexter for his comments on the previous versions of this manuscript. While working on this manuscript (2017-2019), I was supported by a Newton International Fellowship from The Royal Society and by Conicyt PFCHA/Postdoctorado Becas Chile/2017 N° 3140189. Now, I am supported by Fondecyt-Iniciacion 2020 N 1120096, ANID, Chile. Finally, I acknowledge the support from CONICYT PIA APOYO CCTE AFB170008.

## References

Adams J and F Woodward (1989). Patterns in tree species richness as a test of the glacial extinction hypothesis. Nature 339, 699.

Algar AC, JT Kerr, and DJ Currie (2007). A test of metabolic theory as the mechanism underlying broad-scale species-richness gradients. Global Ecology and Biogeography 16, 170–178.

Allen AP, JH Brown, and JF Gillooly (2002). Global biodiversity, biochemical kinetics, and the energetic-equivalence rule. Science 297, 1545–1548.

Allen AP, JF Gillooly, JH Brown, et al. (2007). Recasting the species-energy hypothesis: the different roles of kinetic and potential energy in regulating biodiversity. Scaling biodiversity 1.

Brodie JF (2019). Environmental limits to mammal diversity vary with latitude and global temperature. Ecology Letters 22, 480–485. DOI: 10.1111/ele.13206. eprint: https://onlinelibrary.wiley.com/doi/pdf/10.1111/ele.13206.

Brown JH (2014). Why are there so many species in the tropics? Journal of Biogeography 41, 8–22. DOI: 10.1111/jbi.12228. eprint: https://onlinelibrary.wiley.com/doi/pdf/10.1111/jbi.12228.

Brown JH, JF Gillooly, AP Allen, VM Savage, and GB West (2004). Toward a metabolic theory of ecology. Ecology 85, 1771–1789.

Burrill E, A Wilson, J Turner, S Pugh, J Menlove, G Christiansen, B Conkling, and W David (2018). The Forest Inventory and Analysis Database: database description and user guide version 8.0 for Phase 2. U.S. Department of Agriculture, Forest Service. Available at web address: http://www.fia.fs.fed.us/library/database-documentation.

Cade BS, JW Terrell, and RL Schroeder (1999). Estimating effects of limiting factors with regression quantiles. Ecology 80, 311–323.

Chave J, M Réjou-Méchain, A Búrquez, E Chidumayo, MS Colgan, WB Delitti, A Duque, T Eid, PM Fearnside, RC Goodman, et al. (2014). Improved allometric models to estimate the aboveground biomass of tropical trees. Global change biology 20, 3177–3190.

Cornell HV (2013). Is regional species diversity bounded or unbounded? Biological Reviews 88, 140–165.

Currie DJ (2007). Regional-to-global patterns of biodiversity, and what they have to say about mechanisms. Scaling biodiversity, 258–282.

Currie DJ, GG Mittelbach, HV Cornell, R Field, JF Guégan, BA Hawkins, DM Kaufman, JT Kerr, T Oberdorff, E O’Brien, et al. (2004). Predictions and tests of climate-based hypotheses of broad-scale variation in taxonomic richness. Ecology letters 7, 1121–1134.

Currie DJ (1991). Energy and large-scale patterns of animal-and plant-species richness. The American Naturalist 137, 27–49.

Currie DJ and V Paquin (1987). Large-scale biogeographical patterns of species richness of trees. Nature 329, 326.

Donoghue MJ (2008). A phylogenetic perspective on the distribution of plant diversity. Proceedings of the National Academy of Sciences 105, 11549–11555.

Enquist BJ, JH Brown, and GB West (1998). Allometric scaling of plant energetics and population density. Nature 395, 163–165.

Enquist BJ and KJ Niklas (2001). Invariant scaling relations across tree-dominated communities. Nature 410, 655.

Field R, EM O’Brien, and RJ Whittaker (2005). Global models for predicting woody plant richness from climate: development and evaluation. Ecology 86, 2263–2277.

Francis AP and DJ Currie (2003). A globally consistent richness-climate relationship for angiosperms. The American Naturalist 161, 523–536.

Gaston KJ (2000). Global patterns in biodiversity. Nature 405, 220.

Gillooly JF, JH Brown, GB West, VM Savage, and EL Charnov (2001). Effects of size and temperature on metabolic rate. science 293, 2248–2251.

Hahsler M, C Buchta, B Gruen, and K Hornik (2019). arules: Mining Association Rules and Frequent Itemsets. R package version 1.6–3.

Harris I, TJ Osborn, P Jones, and D Lister (2020). Version 4 of the CRU TS monthly high-resolution gridded multivariate climate dataset. Scientific data 7, 1–18.

Hawkins BA (2001). Ecology’s oldest pattern? Trends in Ecology & Evolution 16, 470.

Hawkins BA, JAF Diniz-Filho, LM Bini, MB Araújo, R Field, J Hortal, JT Kerr, C Rahbek, MÁ Rodríguez, and NJ Sanders (2007). Metabolic Theory and Diversity Gradients: Where Do We Go from Here? Ecology 88, 1898–1902. ISSN: 00129658, 19399170.

Hawkins BA, FS Albuquerque, MB Araujo, J Beck, LM Bini, FJ Cabrero-Sañudo, I Castro-Parga, JAF Diniz-Filho, D Ferrer-Castán, R Field, et al. (2007). A global evaluation of metabolic theory as an explanation for terrestrial species richness gradients. Ecology 88, 1877–1888.

Hawkins BA, EE Porter, and JA Felizola Diniz-Filho (2003). Productivity and history as predictors of the latitudinal diversity gradient of terrestrial birds. Ecology 84, 1608–1623.

Hijmans RJ, SE Cameron, JL Parra, PG Jones, and A Jarvis (2005). Very high resolution interpolated climate surfaces for global land areas. International Journal of Climatology: A Journal of the Royal Meteorological Society 25, 1965–1978.

Jetz W and C Rahbek (2002). Geographic range size and determinants of avian species richness. Science 297, 1548–1551.

Koenker R (2018). quantreg: Quantile Regression. R package version 5.38.

Koenker R and G Bassett Jr (1978). Regression quantiles. Econometrica: journal of the Econometric Society, 33–50.

Koenker R and JA Machado (1999). Goodness of fit and related inference processes for quantile regression. Journal of the american statistical association 94, 1296–1310.

Kreft H and W Jetz (2007). Global patterns and determinants of vascular plant diversity. Proceedings of the National Academy of Sciences 104, 5925–5930.

Magallon S and MJ Sanderson (2001). Absolute diversification rates in angiosperm clades. Evolution 55, 1762–1780.

Maherali H, WT Pockman, and RB Jackson (2004). Adaptive variation in the vulnerability of woody plants to xylem cavitation. Ecology 85, 2184–2199.

Maitner BS, B Boyle, N Casler, R Condit, J Donoghue, SM Durán, D Guaderrama, CE Hinchliff, PM Jørgensen, NJ Kraft, et al. (2018). The bien r package: A tool to access the Botanical Information and Ecology Network (BIEN) database. Methods in Ecology and Evolution 9, 373–379.

Mucina L (2019). Biome: evolution of a crucial ecological and biogeographical concept. New Phytologist 222, 97–114.

Muggeo VM (2008). segmented: an R Package to Fit Regression Models with Broken-Line Relationships. R News 8, 20–25.

Mutke J and W Barthlott (2005). Patterns of vascular plant diversity at continental to global scales. Biologiske skrifter 55, 521–531.

Mutke J, G Kier, G Braun, C Schultz, and W Barthlott (2001). Patterns of African vascular plant diversity: A GIS based analysis. Systematics and Geography of Plants, 1125–1136.

Nee S (2001). Inferring speciation rates from phylogenies. Evolution 55, 661–668.

O’Brien E (1993). Climatic gradients in woody plant species richness: towards an explanation based on an analysis of southern Africa’s woody flora. Journal of Biogeography, 181–198.

O’Brien E(1998). Water-energy dynamics, climate, and prediction of woody plant species richness: an interim general model. Journal of Biogeography 25, 379–398.

Oliveira-Filho A (2014). NeoTropTree, Flora arbórea da Região Neotropical: Um banco de dados envolvendo biogeografia, diversidade e conservação. Belo Horizonte: Universidade Federal de Minas Gerais.

Olson DM, E Dinerstein, ED Wikramanayake, ND Burgess, GV Powell, EC Underwood, JA D’amico, I Itoua, HE Strand, JC Morrison, et al. (2001). Terrestrial Ecoregions of the World: A New Map of Life on EarthA new global map of terrestrial ecoregions provides an innovative tool for conserving biodiversity. BioScience 51, 933–938.

Phillimore AB, RP Freckleton, CDL Orme, and IP Owens (2006). Ecology predicts large-scale patterns of phylogenetic diversification in birds. The American Naturalist 168, 220–229.

R Core Team (2018). R: A Language and Environment for Statistical Computing. R Foundation for Statistical Computing. Vienna, Austria.

Ricklefs RE (2004). A comprehensive framework for global patterns in biodiversity. Ecology letters 7, 1–15.

Rohde K (1992). Latitudinal gradients in species diversity: the search for the primary cause. Oikos, 514–527.

Segovia RA, RT Pennington, TR Baker, FC De Souza, DM Neves, CC Davis, JJ Armesto, AT Olivera-Filho, and KG Dexter (2020). Freezing and water availability structure the evolutionary diversity of trees across the Americas. Science Advances 6, eaaz5373.

Silva de Miranda PL, AT Oliveira-Filho, RT Pennington, DM Neves, Baker, T R, and KG Dexter (AUG 2018). Using tree species inventories to map biomes and assess their climatic overlaps in lowland tropical South America. English. Global ecology and biogeography 27, 899–912. ISSN: 1466-822X. DOI: {10.1111/geb.12749}.

Stephenson NL (1990). Climatic control of vegetation distribution: the role of the water balance. The American Naturalist 135, 649–670.

Wiens J and M Donoghue (DEC 2004). Historical biogeography, ecology and species richness. English. Trends in ecology & evolution 19, 639–644. ISSN: 0169-5347. DOI: {10.1016/j.tree.2004.09.011}.

Wiens JJ (2011). The causes of species richness patterns across space, time, and clades and the role of “ecological limits”. The Quarterly Review of Biology 86, 75–96.

Wright DH (1983). Species-energy theory: an extension of species-area theory. Oikos, 496–506.

Zanne AE, DC Tank, WK Cornwell, JM Eastman, SA Smith, RG FitzJohn, DJ McGlinn, BC O’Meara, AT Moles, PB Reich, DL Royer, DE Soltis, PF Stevens, M Westoby, IJ Wright, L Aarssen, RI Bertin, A Calaminus, R Govaerts, F Hemmings, MR Leishman, J Oleksyn, PS Soltis, NG Swenson, L Warman, and JM Beaulieu (FEB 6 2014). Three keys to the radiation of angiosperms into freezing environments. English. Nature 506, 89+. ISSN: 0028-0836. DOI: {10.1038/nature12872}.

